# Characterization of regeneration initiating cells during *Xenopus laevis* tail regeneration

**DOI:** 10.1101/2023.03.30.534908

**Authors:** Sindelka Radek, Abaffy Pavel, Zucha Daniel, Naraine Ravindra, Kraus Daniel, Netusil Jiri, Smetana Karel, Lukas Lacina, Endaya Berwini Beduya, Neuzil Jiri, Psenicka Martin, Kubista Mikael

## Abstract

Embryos are regeneration and wound healing masters. They not only rapidly close their wounds, remodel injured tissue without a scar, but also regenerate body parts. Many animal models with variable regenerative capabilities have already been studied. Additionally, with the introduction of high throughput techniques, novel regeneration mechanisms including genes and signaling pathways, and specialized cell types required for regeneration control in spatial and temporal aspects have been identified. Until now our knowledge has been limited to primarily the late phases of regeneration (> 1 day post injury). In this paper, we reveal the critical steps for regeneration initiation. We have discovered Regeneration Initiating Cells (RICs) using single cell and spatial transcriptomic analyses during tail regeneration in *Xenopus laevis*. RICs are formed transiently from the basal epidermal cells and are critical for the modification of the surrounding extracellular matrix to allow for migration of other cell types such as regeneration organizing cells that further promote regeneration. Absence or deregulation of RICs leads to excessive extracellular matrix deposition and regeneration defects.

## Introduction

Regeneration is the complete restoration of ‘missing’ tissue with fully functional tissue. It is different from the process of repair, which is associated with scar formation and impaired function[1, 2]. Fishes and amphibians have nearly perfect regenerative capacity during early development, and some show partial regeneration of certain organs like the heart, retina, liver, limb, and kidney even in adulthood[3]. Mammals can regenerate certain tissues such as amputated digit tips and heal wounds only in younghood. This capacity decreases during maturation and is lost in adults except for a few exceptions such as liver regeneration[4], regrowth of skin of spiny mouse[5] and deer antler[6]. Healing capacity declines also with age, leading to age-related disorders such as leg ulcers, diabetic wounds, heart attacks and cerebral infarction[7].

Regeneration is a multi-step process involving many cell types and pathways. Healing is sometimes considered as the initial step of regeneration[8]. Immediately after injury the cell sheets around the wound are activated to prevent further loss of biological material. In fishes and amphibians, the wound epithelium, sometimes referred to as the apical epithelial cap, covers the injured site[9, 10]. This is followed by the transformation of the wound area into a signaling center called the blastema, which comprises specialized cells[11]. The blastema stimulates local cell migration, proliferation and differentiation[12]. This early structure determines the extent of the functional recovery observed in the later phase of tissue regrowth. Many factors are well conserved among vertebrates during the initial step, such as production of small molecules including reactive oxygen species[13], and activation of early response genes followed by the activation of remodeling enzymes[14].

Inflammation burst in the early stages of the regenerative process activates the immune system to protect against infections and stimulates the removal of tissue debris. The role of the immune system during regeneration is multifaceted, and is characterized by the types of immune cells involved and the duration and type of the immune response[15, 16]. Reduced immune response observed in the embryo is one of the factors promoting regeneration in contrast to the strong immune reaction found in adults and phylogenetically advanced organisms, namely mammals[17]. However, the age of the mammal, including humans, can also be associated with different modulations of the inflammatory response, with different ensuing results of healing[18]).

Regeneration continues with cell proliferation and matrix remodeling to ‘fill’ the missing structures, and it results in complete or partial replacement of the injured tissue. Healing results in functionally suboptimal scar formation. In many cases, specialized cell types enter the wound, the scar is organized and the extracellular matrix (ECM), composed mainly of collagen, is established. ECM serves as a provisional scaffold for the remodeling, but it also facilitates cell migration and differentiation during healing. ECM remodeling enzymes have a major role in these processes and are essential for blastema formation[19] and regeneration[20]. The collagen matrix is re-organized and the inflammation is reduced by mechanisms that are not yet fully understood[21]. In organisms with inadequate healing and regeneration, ECM transforms into a fibrous scar that limits tissue remodeling. Many differences in ECM composition between regenerative and repair models have been found and are considered as one of the key factors determining regeneration efficiency[22, 23].

Our understanding of regeneration mechanisms and cellular interactions dramatically changed with the introduction of high throughput methods such as single-cell sequencing[24]. Their application to traditional regenerative models such as the limb of axolotl (*Ambystoma mexicanum)*[25], fin of zebrafish (*Danio rerio*)[26, 27] and digit tip of mouse (*Mus musculus)*[28] has led to the identification of new cell subpopulations required for their regeneration. A breakthrough in the understanding of embryonic regeneration came from the paper studying tail regeneration at the level of individual cells in *Xenopus laevis*[29]. Regeneration organizing cells (ROCs) were discovered and their migration to the injury site and promotion of developmentally related signaling pathways have been observed to stimulate the regrowth of the tail[29]. However, the mechanism of their migration and attraction to the injury site remained unclear. An important advantage of the *Xenopus* model is the presence of a transient refractory stage, where the embryo loses its regenerative capability[16, 30, 31], making it an ideal tool for comparative studies. Comparison of regenerative and refractory stages allowed for the characterization of myeloid cells, which can be divided into either inflammatory or regenerative subpopulations with opposing effects on regeneration. This resembles the difference between mammalian M1 (inflammatory) and M2 (regenerative) macrophages[15].

Single cell studies are however limited by the missing information about cell positions and their surroundings. Spatially resolved transcriptomics is helpful in this context[32]. However, spatial transcriptomic studies of regeneration are scarce, and include recent reports on fibroblast fate during tissue repair[33] and mouse digit regeneration[34]. In our study, we combined results from three high throughput methods: bulk RNA-Seq, single cell RNA-Seq and spatial transcriptomics to investigate regeneration during the initial (hours post amputation - hpa) and later phases (days post amputation – dpa). Our research reveals a new cell type that we refer to as the Regeneration Initiating Cells (RICs). We support the importance of these cells for regeneration through functional studies. This research serves as a novel and detailed study of regeneration initiation, complementing studies in regeneration models that have focused primarily on the later phases of the process.

## Results

### Gene expression during the early states after injury regulates regeneration initiation via a conserved mechanism

We performed bulk RNA-Seq using regenerative and refractory stage embryos to study temporal regulation of gene expression (Fig. 1). In total, 4,358 differentially expressed genes (DEGs) associated with regeneration were identified (adjusted p-value < 0.01; minimal of 20 transcripts in at least one sample). Based on the temporal expression profiles, DEGs were divided into three categories corresponding to the three regeneration (early, middle, late) phases. DEGs expressed during the **early phase** were characterized by an expression burst at 0.5 to 1 hpa followed by a rapid decrease (801 genes, Fig. 2B). Many of these early response genes (e.g. *fos, jun, egr1*) were previously identified in injury models including *Xenopus* embryonic wound healing[14]. Other early phase DEGs coded for ATPases, regulate function of muscle cells (*myh4*, 8, *11* and *13, myl1* and *4, myod1*), oxidative phosphorylation (*ndufa* genes, *cytochrome c oxidase* genes, *sdhc, sdhd* and *ubiquinol-cytochrome c reductase* genes), reactive oxygen species production and additional metabolic changes (catabolic processes). There is a subgroup of early DEGs that is downregulated immediately following the injury (836 genes, Fig. 2B), and are affiliated with GOs associated with muscle activity and ATP metabolic process (Supplement table 1:S2). DEGs with increased expression within 1.5 – 6 hpa (774 genes, Fig. 2B) were classified as the **middle phase**. Their functions are predicted to be associated with tissue remodeling (*mmp1, 8* and *9, timp1* and *3*), cell migration (*epcam, integrin subunits, muc1, vim*) and control of developmental processes (*notch1, shh, wnt10a, sox11, bmp4*). DEGs with increased expression after 1 dpa (1,179 genes, Fig. 2B) are classified to the **late phase**. Their functions are linked to developmental regulation and signaling pathways such as the Wnt (*axin2, dkk1* and *3, lrp1* and *4, ctnnb1, frzb, wnt5b* and *11*) and the TGFβ pathway (*bmp1*, 5 and *7, smad1, 4* and *9, tgfb1* and *2*). Late response was also observed for many *hox* genes, keratins and genes required for proliferation (rRNA metabolism). The complete list of DEGs for each phase and the associated enriched GO terms are available in the Supplement table 1.

**Figure 1.**
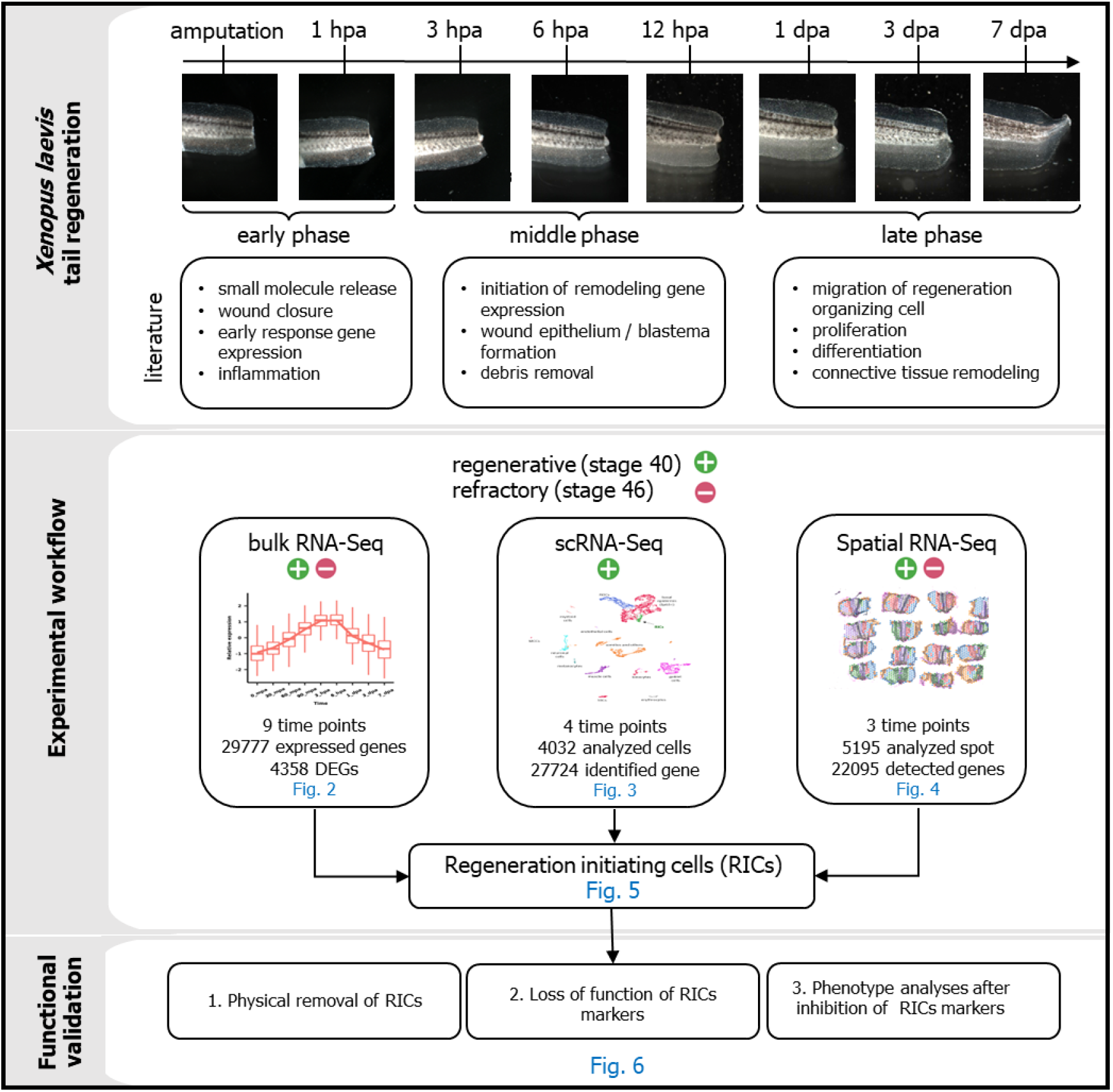
Study design of regeneration initiation using *Xenopus laevis* tail amputation model.

**Figure 2.**
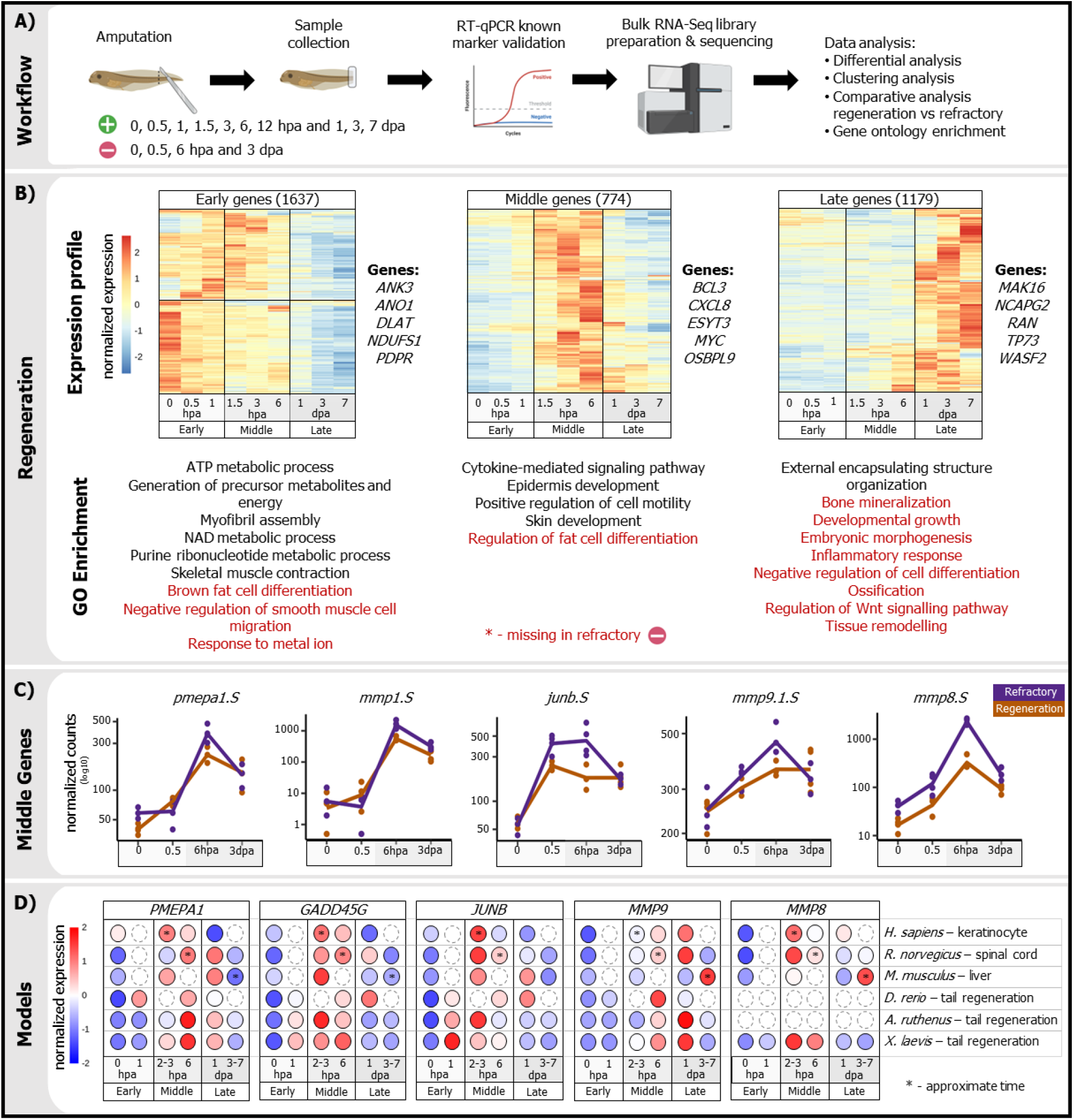
Temporal bulk RNA-Seq comparison between regenerative and refractory embryos. A) Scheme of experiment. B) Regeneration was divided into three phases – early, middle and late. Heatmaps including selected representative genes are showed together with selected enriched GO terms. GO terms in red color are not significantly enriched in refractory samples. C) Five genes from middle group (later identified as RICs markers) were selected for comparison between regenerative (orange) and refractory (purple) samples. D) Targeted analysis of selected middle genes were performed using RT-qPCR and data shows their expression in other regenerative models.

DEG comparison between the regenerative and refractory stage revealed a clear difference during the late phase, with many developmental processes reduced or missing (Fig. 2B, Supplement table 1:S4). This suggests that middle genes determine the consequential regenerative properties of the injured tissue. To see whether the patterning of gene expression after injury is of wider relevance, we analyzed several other regenerative and healing models (scratch assay of human fibroblast layer, and regeneration of rat spinal cord, mouse liver and fish tails) and all showed similar changes in expression of selected middle genes during 3 – 6 hpa, suggesting conservation of regeneration initiation (Fig. 2D). Even though bulk RNA-Seq revealed many potential regenerative candidate genes, characterization of the cell types playing a role during initiation of the process required more detailed single cell analysis.

### Regeneration Initiation Cells (RICs) appear upon initiation of regeneration

We collected and profiled gene expression in 4,032 single cells during the early and middle phases of regeneration with the average gene coverage of ∼6,000 genes per cell (Fig. 1, 3A). RT-qPCR assessment of cell type marker genes between samples extracted from the whole tail versus the cell suspension showed that there were minimal differences in the temporal gene expression profiles due to the disassociation procedure (Supplement Fig. 1). Basic quality controls, such as the level of mitochondrial genes, ribosomal genes and unique molecular identifiers can be found in Supplement Fig. 2. After the clustering and cell type annotation (Fig. 3B, Supplement Fig. 3, 4, Supplement table 2:S1), the basal epidermal cells were found to be the population with the primary temporal changes (Fig. 3B, Supplement Fig. 3). It is from within these basal epidermal cells that we observed a new subpopulation, which we designated Regeneration Initiating Cells (RICs, Fig. 3B). RICs were not present at time 0, which indicates that epidermal cells must first undergo a transition following the amputation. RICs population increased in time, reaching a maximum of 10% of the total basal epidermal cells at 12 hpa (Fig. 3B). Analysis of a previously published single-cell RNA-Seq (scRNA-Seq) experiment that assessed longer time periods of *Xenopus* regeneration, revealed the presence of similar expression signature characteristic of the RICs population[29]. However, in that experiment the RICs were referred to as laminin-rich cells and their function was not further elucidated. Upon re-analysis, we found >50% of overlapping markers between the laminin-rich cells and RICs. In our scRNA-Seq dataset, we identified 272 genes as RICs markers (fold change > 2, *P*_*adj*_ < 0.05), including highest enrichment of *palld*.*L/S, lep*.*L, pmepa1*.*S, inhba*.*L, pthlh*.*S, lamb3*.*L, mmp1*.*S, mmp8*.*L/S* and *lamc2*.*L* (Supplement table 2:S1). Further analysis revealed GO enrichment for processes such as wound healing, cell adhesion, cell migration and ECM remodeling (Fig. 3C, Supplement table 2:S2). The RICs markers significantly (hypergeometric test: pvalue < 0.001) overlapped with the genes from the middle phase of the bulk RNA-Seq (43%), pointing to the contribution of RICs to the middle regeneration phase. Five selected RICs markers with interesting biological relevance were validated using *in situ* hybridization at 1 dpa (Fig. 3D; Supplement Fig. 5). The results showed that these markers are present predominantly in the regeneration bud, indicative of forming an organizational center after injury regeneration onset. However knowledge of cell positions and local cell combination are needed to understand mechanism of the process.

**Figure 3.**
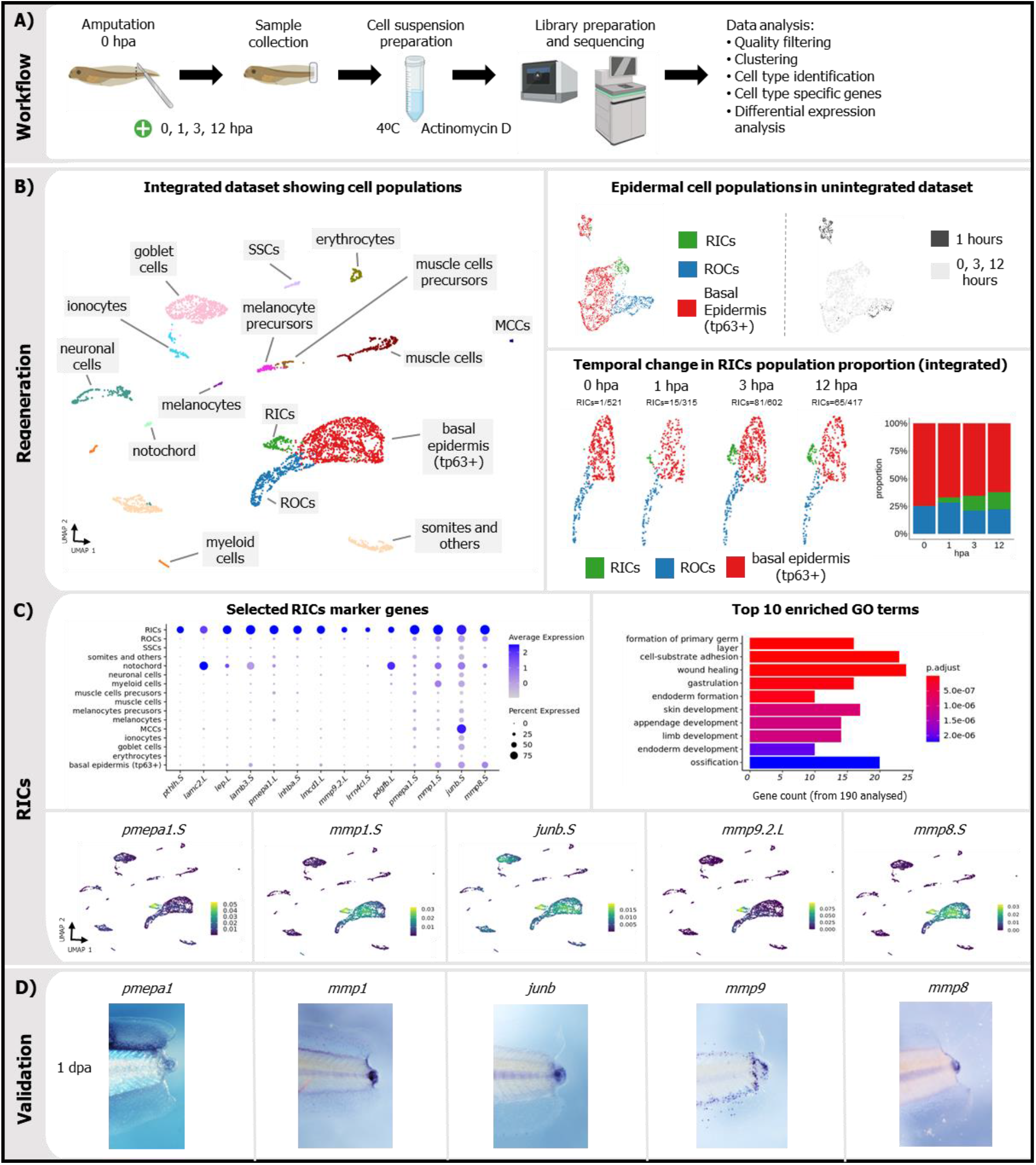
Single cell analysis of regeneration initiation. A) Scheme of scRNA-Seq experiment. B) UMAP visualization of integrated datasets, leading to identification of regeneration initiating cells (RICs), ROCs – regeneration organizing cells, SSCs – small secretory cells, MCCs - multiciliated cells. Temporal changes in epidermal cell populations (tp63+) are shown using unintegrated and integrated results. C) The expression profile of the selected RICs marker genes within the different cell populations. Top ten enriched Gene Ontology terms for the RICs marker genes. D) Validation of RICs markers by *in situ* hybridization at 1 dpa and preferential expression of RIC markers in regeneration bud.

### RICs are accumulated in the regeneration bud during regenerative stage of the embryo

To better understand the spatial arrangement of the RICs population within the regenerating tissue and their involvement during the regeneration and repair processes in the bud after the tail amputation, we performed spatial transcriptomics on embryos in regenerative and refractory stages (Fig. 4A). We assessed times within the middle and late phases. After data quality control (Supplement Fig. 6) and clustering, the majority of cell types characterized in the scRNA-Seq dataset were recovered in the spatial dataset through canonical markers. They include clusters of epidermal, muscle, somite, neural, notochord, myeloid, ROCs and regeneration bud (blastema) markers (Fig. 4B, Supplement table 3:S1).

**Figure 4.**
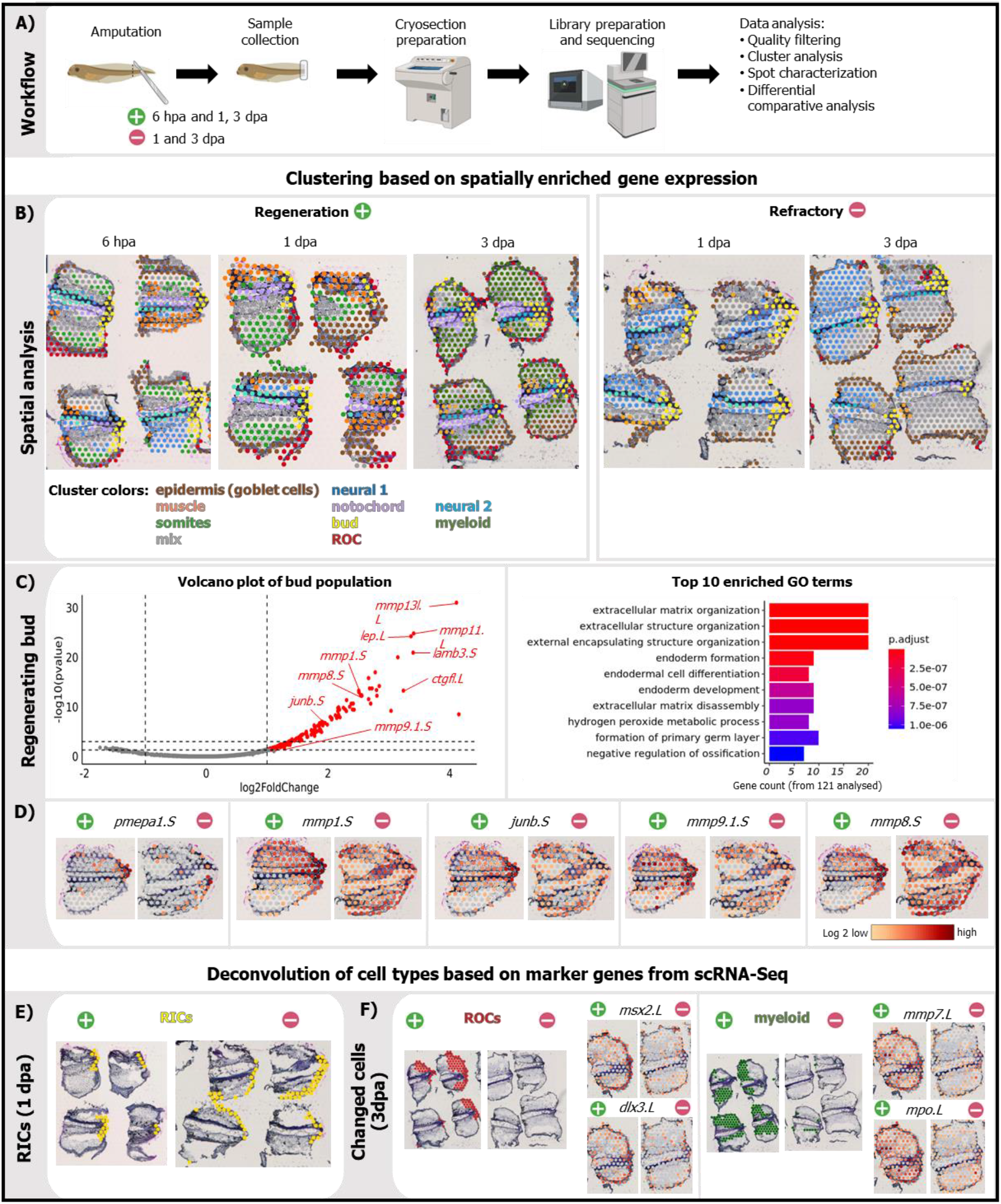
Spatial transcriptomics of tail regeneration. A) Scheme of spatial transcriptomics experiment. B) Clusters formed based on enriched expression in regenerative and refractory samples. Maximum number of 10 clusters were selected in Loupe and manually annotated based on top cluster markers and comparison with scRNA-Seq. C) Analysis of regenerating bud cluster. Markers are shown in the volcano plot while the bar plot shows the top 10 enriched Gene Ontology terms associated these enriched genes. D) Comparison of spatial expression of selected regenerative RICs markers in regenerative and refractory samples at 1 dpa. E) List of RICs markers from scRNA-Seq was visualized in spatial results in regenerative and refractory samples at 1 dpa. F) Visualization of ROCs and myeloid markers (based on scRNA-Seq) in regenerative and refractory samples at 3 dpa.

Results from 6 hpa and 1 dpa of regeneration and refractory stages were analyzed to compare their spatial composition. In the bud ‘spots’ in the regenerative stage, we identified 165 genes to be upregulated (fold-change >2, *P*_*adj*_ < 0.05) when compared with the rest of the tissue. Among them, 46% have been identified as RICs markers in our scRNA-Seq analysis, suggesting a predominant RICs presence in the regeneration bud (Fig. 4C, Fig. 5A). The functions of the regenerative bud genes included ECM organization, collagen degradation, cell differentiation and regulation of developmental processes (Fig. 4C, Supplement table 3:S2). Interestingly, a distinct bud cluster was also identified in the refractory samples. It was characterized by overexpression of 161 genes with 40% of them overlapping with markers of regenerative bud spots (Fig. 5B, Supplement table 4:S1). To visualize differences between regenerative and refractory samples at the individual RICs marker level, we showed that the expression of the marker genes *mmp8*.*L/S, mmp9*.*1*.*S, pmepa1*.*S, mmp1*.*S and junb*.*S* were overabundant in the bud during the regeneration stage, but dispersed throughout the tissue during the refractory period (Fig. 4D).

**Figure 5.**
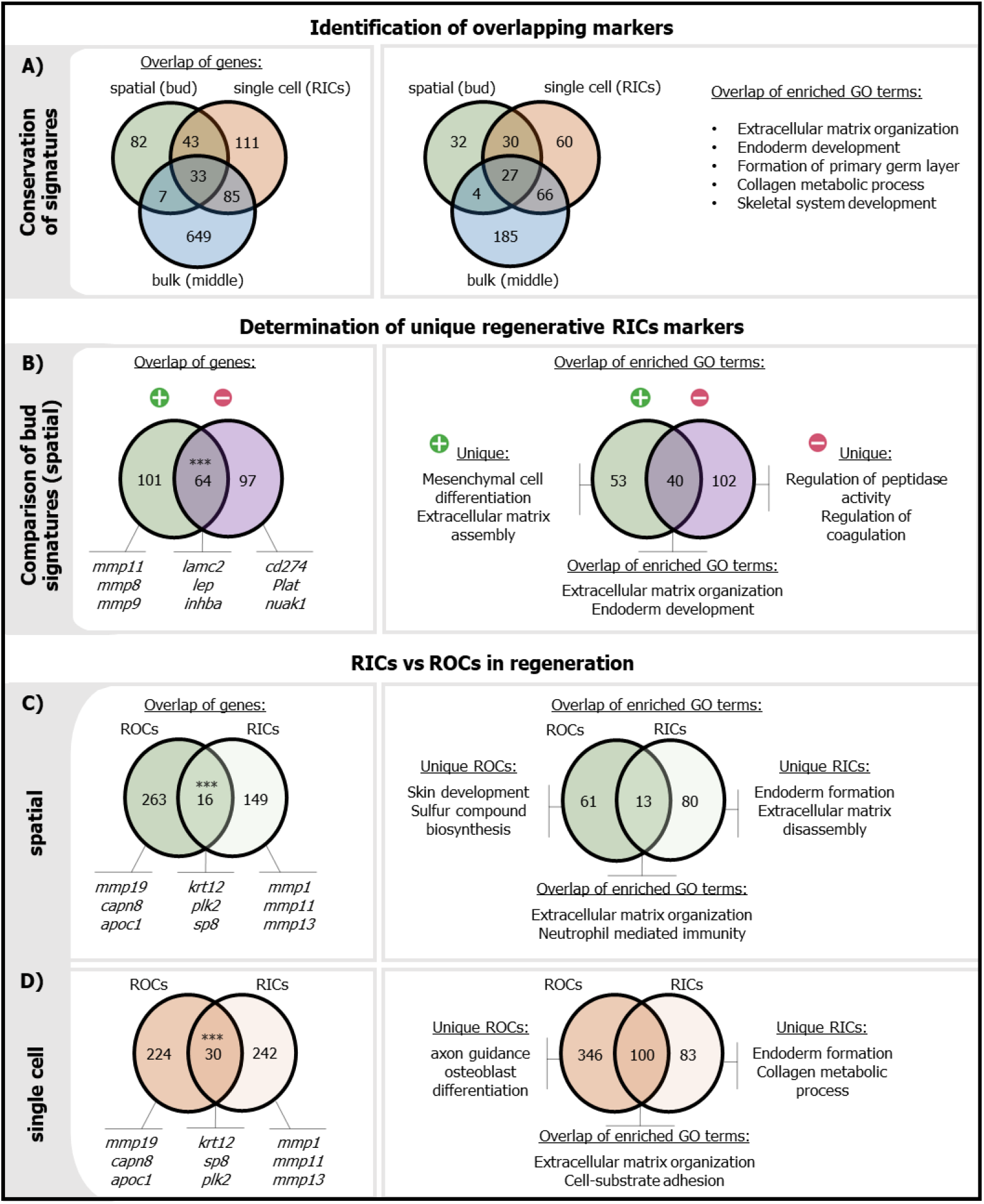
Comparative analysis of descriptive datasets. Overlap of significantly expressed transcripts and their associated enriched Gene ontology terms in the: A) middle bulk RNA-Seq, RICs in single cell and regenerating bud in spatial datasets, B) bud clusters in regenerative and refractory samples in spatial datasets to reveal unique regenerating genes, and C) RICs and ROCs marker genes in spatial and D) single cell datasets to determine their relationship. Significance enrichment of the gene overlap was determined using a hypergeometric test (1-tailed) (ns – not significant, * - P ≤ 0.05, ** - P ≤ 0.01, *** - P ≤ 0.001).

Next, cellular composition of bud spots in the spatial dataset was approximated by deconvolution using our annotated scRNA-Seq dataset as reference. In the regeneration stage, RICs were deconvoluted and found to be primarily localized in the regeneration bud, while in the refractory stage they were largely dispersed on the surface of the embryo at 1 dpa (Fig. 4E). The prominent cell population necessary for regeneration continuation, ROCs[29], were observed to localize to the amputation site of regenerative samples primarily during the later phase at 3 dpa, judged also by spatial plots of canonical ROCs markers (*msx2*.*L* and *dlx3*.*L)*. This spot cluster was completely missing in the refractory samples (Fig. 4F). Marker gene expression of the remaining cell types in the scRNA-Seq dataset (epidermal, muscle, neural and notochord tissues) confirmed their presence in their expected locations (Supplement Fig. 7). Taken together, spatial transcriptomics confirmed the differences of response to injury between the regenerative and refractory embryos suggesting the importance of RICs regeneration initiation.

### RIC marker genes expressed in the middle phase in the bud are essential to initiation of regeneration

In previous parts, we studied and analyzed regeneration initiation independently, using three high-throughput methods: bulk RNA-Seq, scRNA-Seq and spatial transcriptomics. The datasets were further analyzed in order, 1) to study the conservation of identified gene set signatures in time and space; 2) to find regeneration-linked genes using comparison of bud expression signatures in regenerative and refractory samples; and 3) to describe a potential relationship between RICs and ROCs in regenerative samples.

To study the conservation of the responsive gene signatures, the middle category pattern (genes upregulated between 1.5 - 6 hpa; bulk RNA-seq) was compared with single cell RICs markers (scRNA-Seq) and with genes upregulated in bud spots of regenerating embryos (spatial transcriptomics). Overlap of the three sets of genes showed conservation for 33 regeneration-initiating genes (e.g. *mmp1, lep*.*L, mmp8, sox11*.*S*) and 27 shared GOs of biological processes, such as ECM organization and regulation of development (Fig. 5A, Supplement table 4).

In the second comparison, the bud gene expression signatures were compared between regenerating and refractory embryos in the spatial datasets. Out of the 165 markers of the regeneration bud, 64 markers were shared (e.g. *lamc2*.*L, inhba*.*L, fn1*.*S, mmp13*.*L, lep*.*L, gadd45g*.*L*) and 101 were identified as unique to the regeneration bud (Fig. 5B, Supplement table 4:S1). Among the unique RICs markers of regeneration, 20 were conserved within the 33 regeneration-initiating genes (Fig. 5A). This included markers of ECM remodeling (*mmp8, 9* and *11*) and potential regulation of cell differentiation (*rgcc*.*L, sox11*.*S, fibin*.*S*), as was also confirmed on the level of GOs of biological processes (Fig. 5B, Supplement table 4).

In the third analysis, a relationship between RICs and ROCs was studied. Both cell populations are formed from basal epidermal cells (*tp63+*), but they differ in their formation, behavior and gene expression signatures. To better discern the RICs-ROCs relationship, we compared their signatures in spatial and single cell datasets (Fig. 5C, 5D, Supplement table 4). As revealed by the GO analysis, RICs are involved in extracellular space modifications and regulation of development, while ROCs were found to be important regulators of epidermis development, limb and gland morphogenesis and metabolic processes (Fig. 5C). Both ROCs and RICs appear to be important for ECM remodeling, but the ECM related genes they express are different. The ROCs specific genes include ECM components (*emilin1*.*L, nid2 L/S, vcan*.*L*), laminins (*lama5*.*L, lamb1*.*L, lamb2*.*L*), *hyal2*.*S*, keratins (*krt8*.*1*.*L, krt18*.*1*.*L, krt70 L/S*), and collagens (*col14a1 L/S, col18a1 L/S*). They also express several remodeling enzymes such as ADAM metallopeptidase (*adamts10*.*L, adamts12*.*L, adamts18*.*L, adamts7*.*L*) and matrix metalloproteinases (*mmp3*.*L*). In contrast, RICs are rich in expression of integrins (*itga2*.*L, itga5*.*L, itga6*.*S, itgb1*.*L, itgb1*.*S, itgb4*.*L, itgb6*.*L*), laminins (*lama3 L/S, lamc2*.*L)* and matrix metalloproteinases (*mmp1*.*S, mmp8*.*S, mmp9*.*L/S, mmp11*.*L, mmp13l*.*L/S*). Visual inspection of ECM components in scRNA-Seq revealed that the remaining epidermal cells shared with RICs upregulated genes for regeneration-dominant collagens (*col1a1/2, col3a1, col4a5/6, col6a1/2*), and *dsp*.*S, lum*.*L/S, ogn*.*L, dcn*.*L/S, lamb4*.*L*. This suggests that RICs are more similar to the original basal epidermal cells compared to ROCs in regard to ECM. The importance of ECM composition for regeneration is further supported by spatial data at 3 dpa, where many ECM components showed increased levels (e.g. *ogn*.*L, dcn*.*L/S, lum*.*L, hyal4*.*L, vcan*.*L, col1a1*) in regenerative samples compared to the refractory ones (Supplement Fig. 8). Important regulatory control of the activity of MMPs is based on the production of their inhibitors (*timps*). Our single cell analysis showed *timp* expression in different cell types: *timp-1*.*L/S* in RICs together with somite and notochord cells and with expression of *timp2*.*L* in myeloid cells (Supplement Fig. 9). Unique combinations of *mmps* and *timps* may determine different cell function and migration capability.

### Silencing of RICs markers results in regeneration block caused by fibrosis

Here, we identified several genes potentially required for efficient regeneration initiation. To assess their relevance, we performed complementary functional experiments using physical removal of the bud/RICs, inhibition of the top three RICs marker genes (based on cell specificity, position and known relevance for regeneration) and comparison with the phenotype of the refractory stage.

The regeneration bud (containing mainly RICs, indicated by Fig. 4B) was manually removed at 4-6 hpa, and embryos were cultivated in parallel with controls. The removal of the bud resulted in reduced regeneration capability (Fig. 6B), which was caused by extensive fibrosis showed by accumulation of fibronectin resembling scarring in the regenerating area as well as by defect in migration of ROCs indicated by decreased expression of ROCs markers (Fig. 6B). A similar phenotype was observed in embryos with inhibited *mmp8, mmp9* and *pmepa1* activity using Vivo-morpholino oligonucleotides (MOs). Application of MOs for the first day of regeneration resulted in reduced efficacy of the process, accompanied by fibrosis and defects in migration of ROCs (Fig. 6C, D). Refractory samples were prepared for comparison with regenerating embryos, and extensive fibrosis and ROCs migration decrease was also detected here (Fig. 6F). In addition, a migration defect of refractory ROCs was observed in the bulk RNA-Seq and spatial experiments (based on decreased levels of *msx2, dlx3* in refractory compared to regenerating embryos at 3 dpa in spatial and 3 dpa in late phase of bulk RNA-Seq). Altogether, this suggests direct connection between the role of RICs in ECM remodeling and control of ROCs migration.

**Figure 6.**
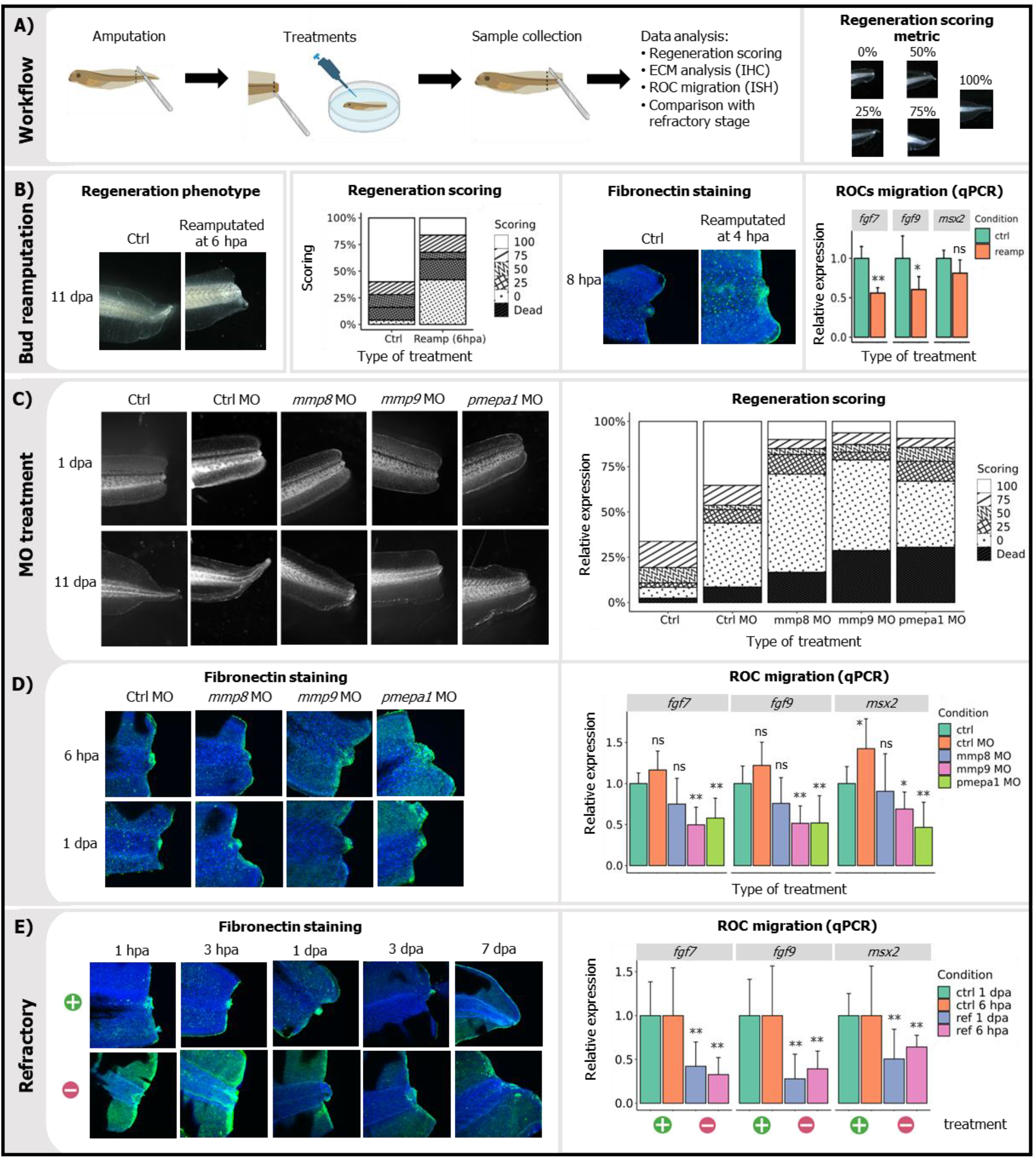
Functional validation of regenerative RICs. A) Scheme of functional experiments. B) Removal of regenerating bud (RICs) at 6hpa and its phenotype (11 dpa), scoring, fibrosis (Fibronectin staining, 8 hpa) and ROC migration defects (RT-qPCR of ROCs markers 1 dpa). C) Loss of function of three RICs markers: pmepa1, mmp8 and mmp9 using Vivo MO. Phenotypes and scoring are shown together with D) extensive fibrosis and ROCs migration defect. E) Extensive fibrosis and ROCs defect in refractory samples. Brightfield and confocal images scale size is 300 µm. RT-qPCR prepared from at least biological triplicates each containing at least five dissected regenerating tails. Significance testing was done using the student T-test (ns – not significant, * - P ≤ 0.05, ** - P ≤ 0.01, *** - P ≤ 0.001).

## Discussion

Here, we propose a novel mechanism of regeneration initiation dependent on the activity of the cell type we refer to as regeneration-initiating cells (RICs). RICs population form from the basal epidermal population and become present at the edge of the amputated tail tissue within a few hoursand feature remodeling of the surrounding ECM to promote the migration of cells required for regeneration. *Xenopus* tail regeneration is dependent on the migration of ROCs, but we can speculate about other potential partners from other models such as connective fibroblasts in axolotl[25, 35], interaction of fibroblasts and keratinocytes in mammalian wound healing[36], or the fibroblast subtype in reindeer antler regeneration[37]. RICs population has not been described prior to this study, because all available single cell studies in the regeneration field have focused primarily on later time points, which resulted in the lack of information about early events of regeneration initiation. Interestingly, our targeted expression analysis showed similar increases in RICs-specific gene expression in early stages post injury in other models of regeneration (including mammals), which indicates an evolutionary conservation of the initiation pattern.

Applying descriptive analyses in combination with functional validation, we found that, interestingly, all three datasets resulted in the identification of a group of genes (putative RICs markers), which are conserved in time and space, and are expressed in a specific cell subpopulation. Comparison between regenerative and refractory stages indicated importance of combination of all three approaches to determine the mechanism of regeneration initiation, especially in the spatial context. Temporal expression profile of RICs markers based on bulk RNA-Seq data is characterized by their rapid increase during 3-6 hpa, followed by a continual decline of expression after 1 dpa. A burst of early response gene expression right before the RICs appearance, suggests a potential link between them. Several studies have confirmed a dependence of the remodeling enzyme expression on the early response genes[38, 39]. As shown by our datasets, RICs are not present in the developing embryos, but are formed following the injury in contrast to ROCs, which are present at the tail growing tip[29]. RICs and ROCs share their origin from the basal epidermal cells, but RICs are newly formed at the amputated plane, and they later accumulate in the regeneration bud. The basal position of RICs and ROCs in the epithelium can be a general feature in vertebrates, because proliferating keratinocytes capable of the wound closure are located in the basal layer of epidermis in mammals, and this can be employed in wound therapy in the clinic[40]. No sign of active migration of RICs was observed in our spatial analysis, but cell migration is required for other cell types (ROCs[29] and myeloid cells[15, 41]) in order to regulate regeneration. We hypothesized that the role of RICs during regeneration is linked to stimulation of cell migration (e.g that of ROCs) based on their markers, including enriched ECM remodeling enzymes. However other migratory cells using alternative routes, such as primitive myeloid cells, were not affected by silencing of RICs (Supplement Fig. 10).

Based on our single cell results, RICs specifically express more than a hundred genes with various functions. There is evidence in the literature about requirement of these genes for regeneration. For example, *junb* was identified in *Xenopus* tail regeneration, and it has a role in the regulation of cell proliferation following induction by TGFβ signaling[42]; the PMEPA1/TMEPA1 protein coded by the gene *pmepa1* acts as a negative regulator of TGFβ signaling, reducing cell migration[43]. There are also contradictory results showing promoting of malignant cell migration by PMEPA1/TGFβ[44], suggesting the well-documented context dependency of the TGFβ signaling in cancer. In addition, RICs also expressed *tgfbi* and *inhba*, which are members of the TGFβ family regulating cell adhesion[45] and modulating TGFβ pathway activity[46]. However, whether expressed TGFβ members in *Xenopus* RICs act positively or negatively with regards to TGFβ signaling cannot be deduced from our results. The homeolog of another RICs marker (*gadd45g*) Gadd45 is involved in effective wound healing of *Drosophila* and protects cells from DNA damage during high generation of ROS [47]. Gadd45b was also found to be required for liver regeneration[48].

One of the most dominant RICs markers are genes coding for enzymes remodeling the extracellular matrix, especially *mmps*. There is extensive literature about the function of *mmps*/MMPs, their substrates and activities[49, 50]. MMPs are enzymes with multiple functions, expressed by a variety of cell types, and can be found active in normal and pathological situations[51-54]. However, the numerous members of the MMP family makes detailed experimental characterization challenging[55]. The most important function of MMPs is the cleavage of ECM components which then leads to the subsequent changes to the behavior of the neighboring cells. However, cleavage of ECM could also lead to signaling stimulation. Signaling role of MMPs was suggested in the context of cleavage of laminin 111 to regulate the epithelial-mesenchymal transition[56], laminin 5 gamma 2 chain to induce cell migration[57], and E-cadherin to facilitate migration of epithelial cells[58]. The importance of MMPs during regeneration and healing is supported also in various animal models. Loss of *mmp9* function in zebrafish resulted in defects during wound healing and regeneration via excessive collagen formation [59]. In addition, *mmps* and their inhibitors were identified in zebrafish fin regeneration[60], and several *mmps* including *mmp9* were also induced during axolotl limb regeneration[61]. *MMPs 1, 2, 3, 9, 10, 11, and 13* are highly expressed during the early phase of wound healing in humans after laser treatment[62]. *MMPs 1, 8, and 13* are active as anti-fibrotic and proliferative factors during mouse liver regeneration[63]. Increased expression of several *mmps* (8, 11, 13) was shown also in axolotl and lungfish late phase limb and fin regeneration[64]. Also of relevance, cooperation of individual *mmps* has been suggested during wound healing, which includes the RICs markers *MMP8* and *MMP9*[65].

ECM composition is one of the key factors determining cell migration and behavior, and both RICs and ROCs express many ECM components. Interestingly, this expression pattern is rather exclusive, suggesting production of two types of ECM leading to different cell phenotypes. This implies the future importance of a detailed ECM structure analysis during regeneration for understanding of the mechanism.

Functional experiments revealed the importance of RICs and expression of their markers for regeneration initiation. The removal of the bud tip or silencing of the RICs markers showed phenotypes that were very similar to the refractory stage embryos, which leads us to speculate that RICs accumulated in the bud are required for ECM remodeling following amputation and they are also required for generation of cell migration stimuli. We can also propose the parallel possibility that the regeneration bud acts as a signaling center producing chemoattractants for migratory cells. However, the nature of these putative attractants remains unclear. Of note, several upregulated genes identified in RICs, namely *itga2, lamb3*, and *lamc2* were recently identified as critical molecules predicting an aggressive course in pancreatic cancer[66]. This suggests the importance of such genetically conserved machinery for pancreatic cancer cells and their potent regulatory role in the context of the tumor microenvironment. Linked to this, pancreatic cancer is frequently associated with extensively developed stromal component. Further, the extent of fibroplasia resembles scar tissue[67].

There are studies supporting the importance of the initial step of regeneration (within hours after the injury) as a determinant of regeneration capability. Limb regeneration in adult *Xenopus* is minimal following amputation, but it can be greatly improved by treatment of the stump with pro-regenerative multidrug mixtures only for the first day after injury, resulting in 18 months regeneration. Relevant to this is the applications of two compounds: 1,4-DPCA (1,4-dihydrophenonthrolin-4-one-3-Carboxylic acid), which reduces excessive collagen deposition and resolvin D5 which have anti-inflammatory activity [68]. A recent pre-print showed the role of an *mmp-1* homeolog in the regeneration of the planarian flatworm, *Schmidtea mediterranea* [69], which leads us to speculate about similarities of regeneration regulation also in invertebrates. In summary, we have described a novel cell type involved in regeneration initiation, and we have identified several putative targets for treatment in case of poor healing and regeneration capabilities. There are similarities of the expression profiles we present here to data from prenatal development, wound repair and cancer [70]. This is exemplified by the production of ECM and its remodeling, which is important for both wound healing and cancer spread[71, 72]. We propose that effective therapy will not be based on a single factor, but rather on a combination of several factors applied during individual phases of regeneration.

## Methods

### Embryo preparation and amputation

*Xenopus laevis* females were stimulated with 500 U of human chorionic gonadotropin (Sigma-Aldrich). Eggs were collected the following day and fertilized by testes suspension which were surgically obtained from the male. After the removal of jelly coats by the 2% cysteine treatment, embryos were incubated in 0.1x MBS until the experimental procedure. Amputation was performed manually (removal of ∼30% of tail tissue) using tricaine anesthetized tadpoles at stage 40 (regenerative) and stage 46 (refractory). Embryos were immediately transferred to the 0.1x MMR solution with gentamycin. Solution was changed every day. Embryos for RNA work were again anesthetized at defined time points and the regenerating tissue was dissected, collected and 1) stored at -80°C (bulk RNA-seq, RT-qPCR), 2) dissociated into cell suspension (scRNA-Seq) or 3) embedded to cutting medium (spatial transcriptomics). Whole embryos were fixed in 4% paraformaldehyde for immunohistochemistry or *in situ* hybridization.

### RT-qPCR

Regeneration/refractory tissues were pooled from three embryos and in at least biological triplicates and stored at -80°C. Regenerating tissues were prepared also from tail in sturgeon embryo, spinal cord in rat, fibroblasts from human, liver in mouse and tail in zebrafish embryo (at least biological triplicates). Total RNA was extracted using TriReagent extraction and LiCl precipitation (Sigma) according to the manufacturer’s instructions. The concentration of total RNA was determined using a spectrophotometer (Nanodrop 2000; ThermoFisher Scientific), and the quality of RNA was assessed using a Fragment Analyzer (Agilent, Standard Sensitivity RNA analysis kit, DNF-471).

The cDNA was prepared using 100 ng of total RNA, 0.5 μl of oligo dT and random hexamers (50 μM each), 0.5 μl of dNTPs (10 mM each), 0.5 μl of Maxima H Minus Reverse Transcriptase (ThermoFischer Scientific), 0.5 μl of recombinant ribonuclease inhibitor (RNaseOUT, Invitrogen), 2 μl of 5 × Maxima RTA buffer (Thermo Scientific), which were mixed with Ultrapure water (Invitrogen) to a final volume 10 μl. Samples were incubated for 5 min at 65°C, followed by 10 min at 4°C, 10 min at 25°C, 30 min at 50°C, 5 min at 85°C and cooling to 4°C. Obtained cDNA samples were diluted to a final volume of 60 μl and stored at -20°C.

qPCR reaction contained 5 μl of TATAA SYBR Grand Master Mix, 0.5 μl of forward and reverse primers mix (mixture 1:1, 10 μl each), 2 μl of cDNA and 2.5 μl of RNase-free water. qPCR was performed using the CFX384 Real-Time system (BioRad) with conditions: initial denaturation at 95°C for 3 min, 40 repeats of denaturation at 95°C for 10 s, annealing at 60°C for 20 s and elongation at 72°C for 20 s. Melting curve analysis was performed after to test reaction specificity and only one product was detected for all assays.

### Bulk RNA-Seq

Bulk RNASeq analysis was done to analyze the temporal changes in gene expression during the regeneration and refractory periods. First regeneration experiment assessed time intervals (0, 0.5, 1.5, 3, 6 hpa and 1, 3, 7dpa), while the second which was run in parallel with the refractory, analyzed time intervals (0, 0.5, 6 hpa and 3dpa). The first experiment was done in triplicates while the second used four replicates. All samples were stored at -80°C. Total RNA was extracted from 20 embryos per sample at regenerative and refractory stages. Total RNA was extracted and validated using the same protocol as for RT-qPCR. 100 ng of total RNA was used for library preparation (Lexogen SENSE Total RNA-Seq Library Prep Kit). Library quality was tested by capillary electrophoresis on Fragment Analyzer (Agilent, NGS High Sensitivity kit DNF-474). The libraries were pooled and sequenced using Illumina NextSeq 500, 2×76 bp. On average, approximately 4.6M reads per sample were obtained after quality control. Reads were filtered for low quality reads and adaptor sequences, using TrimmomaticPE (v. 0.36)[73], while ribosomal RNA reads were filtered out using Sortmerna (v. 2.1b)[74] (default parameters). The cleaned reads were then aligned using STAR (v. 2.7.9a)[75] to the *Xenopus laevis* genome version 10.1 and annotation version XENLA_10.1_GCF[76]. A count table was then generated using the python script htseq-count (v. 0.6.1p1)[77] with the parameter “ –m union”. The counts were normalized and analyzed using DESeq2 (v. 1.32.0)[78] under R (v. 4.1.0)[79]. Outlier samples were assessed by analyzing the PCA of the 500 most variable genes. The differential gene expression (DEG) was assessed across the time point for each separate experiment, using the Design: ∼Replicate + Condition, Test: LRT and Reduced model: ∼Replicate. A DEG was defined by a p-adjusted value < 0.01 and with a transcript count of >20 in at least one sample. Clustering to identify DEGs with similar temporal profiles was then done using the degPatterns function from DEGreport (v. 1.28.0)[80]. Parameters for the clustering were restricted to DEGs that had a reproducible minimum of 1.5-fold (short time period) or 2 fold (long time period) difference between any given time point. The resulting clusters were then manually curated to create three final clusters representing early genes (highest expression found between 0 – 1.5 hpa, middle genes with highest expression between 3 -6 hpa and late gene with highest expression between 1 – 7 dpa. Gene ontology (GO) enrichment for each cluster was analyzed using EnrichGO from ClusterProfiler (v. 4.0.5)[81], with the background genes set as the annotatable genes from *X. laevis* (v. 10.1) as reference, but the GO terms from the reference human database org.Hs.eg.db (v. 3.13.0)[82]. GO terms were deemed as significantly enriched using a p-adjusted value of 0.01 after multiple hypothesis correction using Benjamini-Hochberg method. The DEGs and GO terms were then compared between the similar clusters of the different conditions to identify for similarities and differences.

### Single cell analysis (scRNA-Seq)

Single cell experiment was performed using 0, 1, 3, 12 hpa of *Xenopus* tail at stage 42. To avoid artificial activation of gene expression and keep cells at low temperature, whole tissue dissociation process was done in refrigerated room at 4 °C. Embryos were anesthetized and the regenerating part was dissected and collected into 0.5 ml of 2/3 PBS (Sigma D8537, diluted by RNase-free water) with Actinomycin D (ActD, 50 μg/ml; Sigma A1410, storage solution 5 mg/ml in DMSO). Tail pieces from 50 embryos were collected per one sample. Using a test experiment, the artificial expression or loss of cell types within the cell suspension relative to the collected regenerating tissue were assessed using RT-qPCR of known early response genes and cell population markers.

Tubes were spin down for 5 s using a table centrifuge, the medium removed and tissue resuspended in 200 μl of dissociation solution I containing (papain (SERANA, RPL-001-100ml), BSA (40 μg/ml, Thermofisher AM2616), ActD (50 μg/ml), protease (0.5 mg/ml; Sigma P5380, storage solution 250 mg/ml in PBS) and DNase I (50 μg/ml; Roche 11284932001, storage solution 10 mg/ml in RNase-free water)). Dissociation was gently resuspended using P200 wide-bore tip for one minute and gently rotated for 2 minutes. After three repeated resuspending/rotating steps, samples were shortly spun down, supernatant with cells were collected into tubes with 1 ml of fetal bovine serum (FBS, Gibco) prechilled on ice. Undissociated pieces were resupended in another 200 μl of dissociation solution I and dissociated for another three steps of resuspending and rotating. Supernatant with cells was collected into new tube with 1 mL of FBS. Next, tissue was resuspended in 200 μl of dissociation solution II (papain with CaCl_2_ (5 mM), BSA (40 μg/ml), ActD (50 μg/ml), protease (5 mg/ml) and DNase I (50 μg/ml)) and dissociation was performed for three rounds of 10 minutes dissociation the same as described above. In total, five tubes with single-cell suspension in FBS were collected per sample. This allowed for both the preservation of the quality of sensitive cells (released first) and also the collection of the inner cells (collected last).

All tubes were centrifuged (300 g, 6 minutes, 2°C, minimal acceleration, no break), medium was removed (minimal volume was retained to avoid air contact with cells), and cell pellets from one sample were resuspended using P1000 wide-bore tip and pooled together into one 1.5 ml centrifugation tube using 500 μl of 2/3 PBS with BSA (40 μg/ml) and ActD (5 μg/ml). Cells were washed twice using centrifugation (100 g, 7 minutes, 2 °C, minimal acceleration, no break) and resuspended in 1 ml of 2/3 PBS-/-with BSA and ActD. After the third centrifugation (100 g, 7 minutes, 2 °C, minimal acceleration, no break), cell pellet were resuspended in 200 μl of 2/3 PBS with BSA and without ActD. Cell suspensions were filtered using 50 μm filter (CellTricks) and cell concentrations were measured using TC20 Automated Cell Counter (BioRad). Cells larger than 7 μm were counted and samples with viability higher than 80% were used in the next step.

Sequencing libraries were prepared according to the manufacturer’s manual “ Chromium Single Cell 3’ Reagent Kits User Guide (v 3.1)”. In brief, in total 2400 cells per sample were loaded into Chromium chip. Library quality was tested by capillary electrophoresis on Fragment Analyzer (Agilent, NGS High Sensitivity kit, DNF-474). The sample libraries were then pooled and sequenced on Illumina NovaSeq 2000 targeting 100000 read pairs per cell.

Data were processed using STAR v2.7.9a[75]. Reads were aligned and counted against *Xenopus laevis* genome (v 10.1) with annotation XENLA_10.1_GCF obtained from Xenbase[76]. STAR parameters were set “ --soloType CB_UMI_Simple --soloCBwhitelist 3M-february-2018.txt --soloCBstart 1 –soloCBlen 16 --soloUMIstart 17 --soloUMIlen 12 --soloBarcodeReadLength 0 --soloFeatures Gene -- soloCBmatchWLtype 1MM_multi_Nbase_pseudocounts --soloUMIdedup 1MM_Directional_UMItools -- soloUMIfiltering MultiGeneUMI --clipAdapterType CellRanger4 --sjdbGTFfeatureExon exon -- sjdbGTFtagExonParentTranscript transcript_id --sjdbGTFtagExonParentGene gene_name --sjdbOverhang 119”. Raw unfiltered data were processed using R packages. Firstly, droplets with cells were selected using DropletUtils v1.14.1 (20, 21) using command emptyDrops with parameter “ lower = 1000” and FDR <= 0.001. Filtered matrix were used for next processing using Seurat v4.1.0[83]. The normalization and integration of the data followed the standard Seurat protocol. Cells were kept that had less than 15% reads from mitochondrial genes and number of UMIs and counts greater than 2500. The counts were normalized using SCTransform, after which the normalized data from all time points were integrated using integration anchor points selected from the most 10000 variable genes. Identification of nearest neighbors and clusters were done using a resolution of 0.5 and 1:25 PCAs. Uniform manifold approximation and projection (UMAP) was then used to visualize these clusters in a two-dimensional space. Clusters that showed mixed annotation were removed from the final UMAP visualization. FindAllMarkers function was then used to identify marker genes for each cluster using a min.pct of 0.25.

### Spatial transcriptomics

Because of the small size of *X. laevis* tails, biological replicates (6 for refractory and 8 for regenerating embryos) for each condition (Fig. 4A) were prepared and analyzed using the Loupe software. Together we analyzed 1) regenerative 6 hpa and 1 dpa, 2) refractory 6 hpa and 1 dpa, 3) regenerative and refractory 3 dpa). Dissected tissues were embedded and oriented in 50% optimal cutting temperature (OCT) medium, rapidly frozen using dry ice and then transferred to -80°C for a maximum of six weeks of storage. Samples were sectioned sagitally (20 µm thickness) using Leica CM1950 cryostat (Leica Microsystems). Sections collected for the 10X Visium Spatial Gene Expression processing were then stored at -80°C.

Fixation, staining and imaging was performed strictly according to manufacturer’s manuals as in “ Methanol fixation + H&E Staining Demonstrated protocol” (CG000160) and “ Imaging Guidelines Technical Note” (CG000241). In brief, the sections were shortly incubated to thaw them, after which they were methanol-fixed, isopropanol-incubated, and H&E stained. Stained sections were imaged using Carl Zeiss AxioZoomV16 upright microscope equipped with Plan-Neofluar Z objective (2,3x magnification, 0,57 NA, 10,6 WD) at total 63,0x zoom. Samples were imaged with 5% overlap and stitched using ZEN blue pro 2012 software.

Sequencing libraries were prepared according to the manufacturer’s manual “ Visium Spatial Gene Expression Reagent Kits User Guide” (CG000239). Sections were permeabilized for 9 minutes, as precedingly determined by performing optimization experiment. Reverse transcription was performed on the slide, followed by 2nd cDNA strand synthesis. Double-stranded cDNA was transferred, PCR-amplified, enzymatically fragmented, size selected and tagged by Illumina sequencing adapters. Library quality was tested by capillary electrophoresis on Fragment Analyzer (Agilent, NGS High Sensitivity kit DNF-474). The sample libraries were then pooled and sequenced on Illumina NovaSeq 2000 targeting 50000 read pairs per spot. We selected the clearest four neighboring biological replicates for visualization in Fig. 4. In total 5195 spots were analyzed and in average 3800 genes per spot were identified (in total 22095 genes were detected). Ten spot clusters were created (K-means) and annotated based on known predominant marker genes and confirmed using overlap with the single cell data.

For deconvolution of spatial data using single cell results, the raw sequencing data were processed using the recommended set of Space ranger function (v1.2.2, 10x Genomics) for processing of fresh frozen samples. Binary base call files were demultiplexed using mkfastq function with default parameters. The resulting fastq files were mapped separately to Space ranger reference (XENLA_10.1_GCF) using count function which takes a microscope slide image and fastq files, performs alignment, tissue detection, fiducial detection, and barcode/UMI counting. Quality control was done using scanpy 1.9.1 package under python 3.8. Spots were kept, that had less than 20% reads of mitochondrial genes. The data were then log-normalized with a scale factor of 10,000 followed by scaling of data using scanpy package. Then highly variable genes were selected using the built-in function in scanpy. Principal Component Analysis followed by calculation of the nearest neighbors was performed using scanpy built-in functions. Leiden algorithm was used for clustering. UMAP visualization of clusters was performed with built-in function of scanpy package. Cell type identification for each cluster was performed based on known marker genes from single cell RNA-seq. Deconvolution was performed using stereoscope 0.3. For gene selection the top DEGs per cluster were used with minimum fold change set to 1. Training of the model was performed with default parameters and max epochs set to 3000.

### Comparative analysis

The marker genes *(PMEPA, GADD45G, JUNB, MMP9, MMP8*) for the RICs population was assessed in other healing/regenerating tissues from other models to determine if there was also a peak expression during the 2-6 hpa period. The following temporal RT-qPCR experiments were assessed in regenerating: 1) tail in sturgeon, spinal cord in rat, fibroblasts from human, liver in mouse and tail in zebrafish. Values were normalized against the time 0 and presented as relative quantities.

Comparison of the shared genes between the markers for RICs And ROCs population in the single cell, spatial and the middle genes from the bulk RNA-Seq was done. ROCs and RICs markers were filtered to only include those from the scRNA-Seq and spatial experiment that had p-adjusted values < 0.05 and fold change > 2. The enriched GO for these populations were compared between the experiments and also different cell populations using the same method from EnrichGO as described earlier.

*Xenopus laevis* genome (v 10.1) with annotation XENLA_10.1_GCF[76] was used for all alignments and annotations. The unannotated protein coding transcripts were analyzed for their closest ortholog relative to the *H. sapiens* proteome (GRCh38.p14) using the reciprocal best alignment heuristic tool Proteinortho (v. 6.0.31) along with DIAMOND (v. 2.0.11)[84, 85].

### Functional experiments

Reamputation was performed using manual dissection of regeneration bud containing RICs by scalpel at 6 hpa and embryos were incubated in 0.1x MMR with gentamycin at 16°C. Scoring was performed at 7-10 dpa. It was based on three main criteria: notochord length, notochord shape and epidermis attachment to the notochord and scaled from 0 (no regeneration) to 100 (perfect regeneration), samples for Fibronectin staining and RT-qPCR of ROC markers were collected at 1 dpa.

Three Vivo morpholino antisense oligos (MOs) were designed and ordered from Gene Tools, LLC (Philomath, Oregon USA). MOs targeted exon – intron junction near the start of coding sequence of *mmp8, mmp9* and *pmepa1* genes (*mmp8* MO: 5′-ATGAAAACCAATCTACTTACCTCAG-3’; *mmp9* MO: 5′-CATGATCAATAATCCCCTCAC-3’; pmepa1 MO: 5′-CCTTTATAATTGCTACTTACAGATC -3’). Stock solution with concentration 0.5 mM was prepared in UltraPure DNase/RNase free water (Invitrogen) and stored at room temperature. Working solution was diluted using 0.1x MMR with gentamycin resulting in a final 250 nM concentration. Embryos were transferred to MO solutions immediately after tail amputation and incubated for 1 dpa (RT-qPCR, Fibronectin staining) or 7-10 dpa (regeneration scoring). Refractory embryos were prepared the same way as MOs.

Embryos for *in situ* hybridization and immunohistochemistry were fixed in 4% paraformaldehyde overnight at 4°C and then stored in 100% methanol (Penta chemicals) at -20°C. Whole mount *in situ* hybridization protocol was adopted from Sive et al. 2000[86]. At least seven embryos per condition were prepared. Pictures were taken using Nikon SMZ 1500 microscope. Fibronectin immunohistochemistry was performed using at least five embryos per condition. After three washes in PBT for 15 minutes, the primary antibody (anti-Fibronectin, Sigma F3648) was added in concentration 1:150 for overnight incubation at 4°C. The next day secondary antibody was added in concentration 1:500 and incubated overnight at 4°C. DAPI 1:1000 was used as reference nuclei label.

## Acknowledgemets

This work was funded by 86652036 from RVO, the Grant Agency of the Czech Republic GA17-24441S, Operational Programme Research, Development, and Education within the projects: Centre for Tumour Ecology—Research of the Cancer Microenvironment Supporting Cancer Growth and Spread (reg. No. CZ.02.1.01/0.0/0.0/16_019/0000785), project National Institute for Cancer Research (Programme EXCELES, ID Project No. LX22NPO5102) - funded by the Euro-pean Union - Next Generation EU, and by Charles University project Cooperatio ONCO.

