## Supplement Figures for "Characterization of regeneration initiating cells during *Xenopus laevis* tail regeneration"

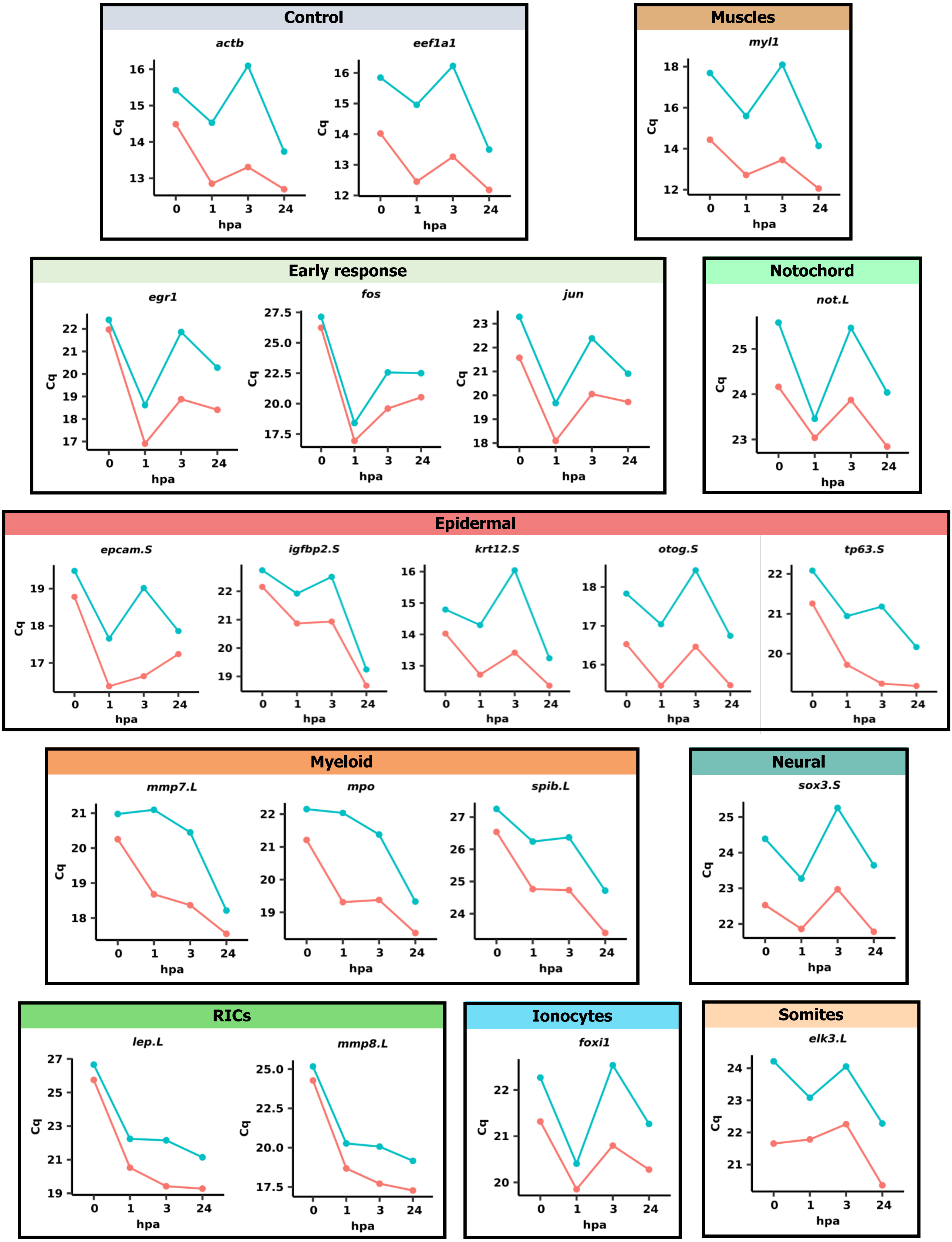


**Supplement Figure 1.** qPCR verification of the temporal effects of the disassociation process on different cell populations. Selected marker genes for the cell populations were assessed in RNA extracted from the tail tissue versus those from cell suspensions. Similarities in the temporal profiles indicated that there was limited effect from the disassociation process.

**Key**
Tails
Suspension

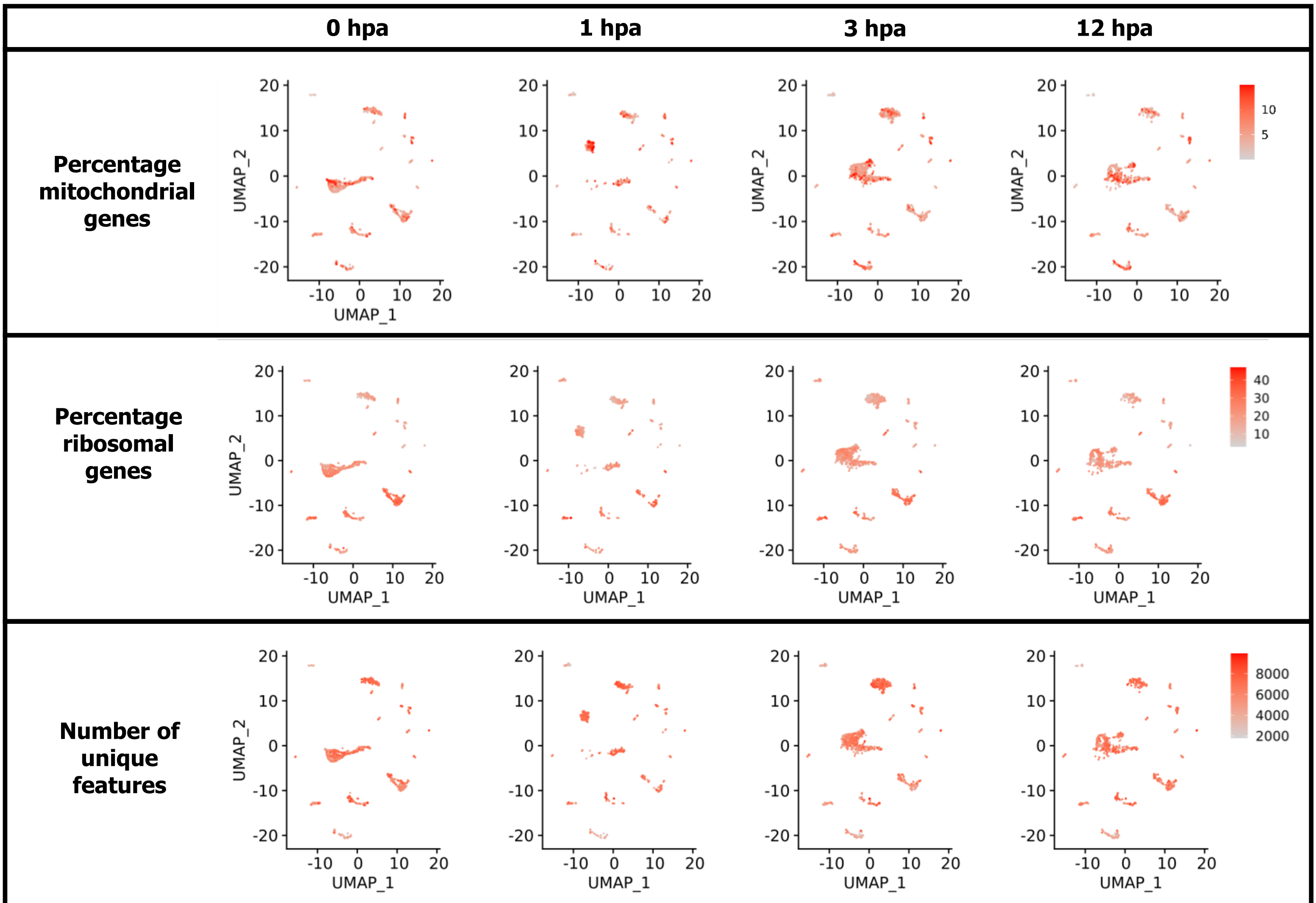


**Supplement Figure 2.** UMAP of unintegrated dataset showing the distribution of the percentage of the mitochondrial and ribosomal genes in each cell, and also the distribution of the number of unique features in each cell after quality control filtering.

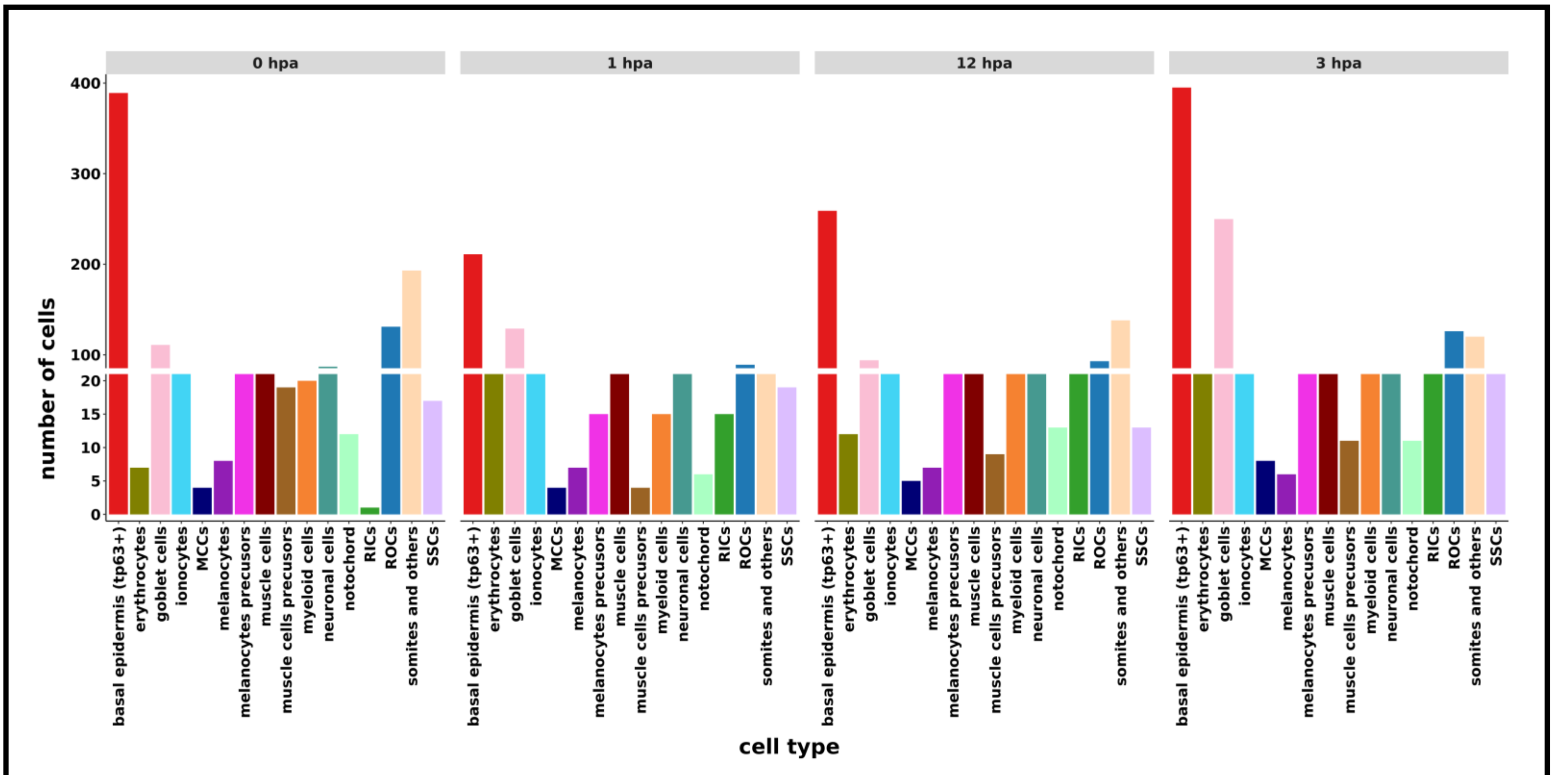


**Supplement Figure 3.** Number of annotated cells identified at each time point in the scRNA-Seq experiment for the regenerating *Xenopus laevis* tail tissue.

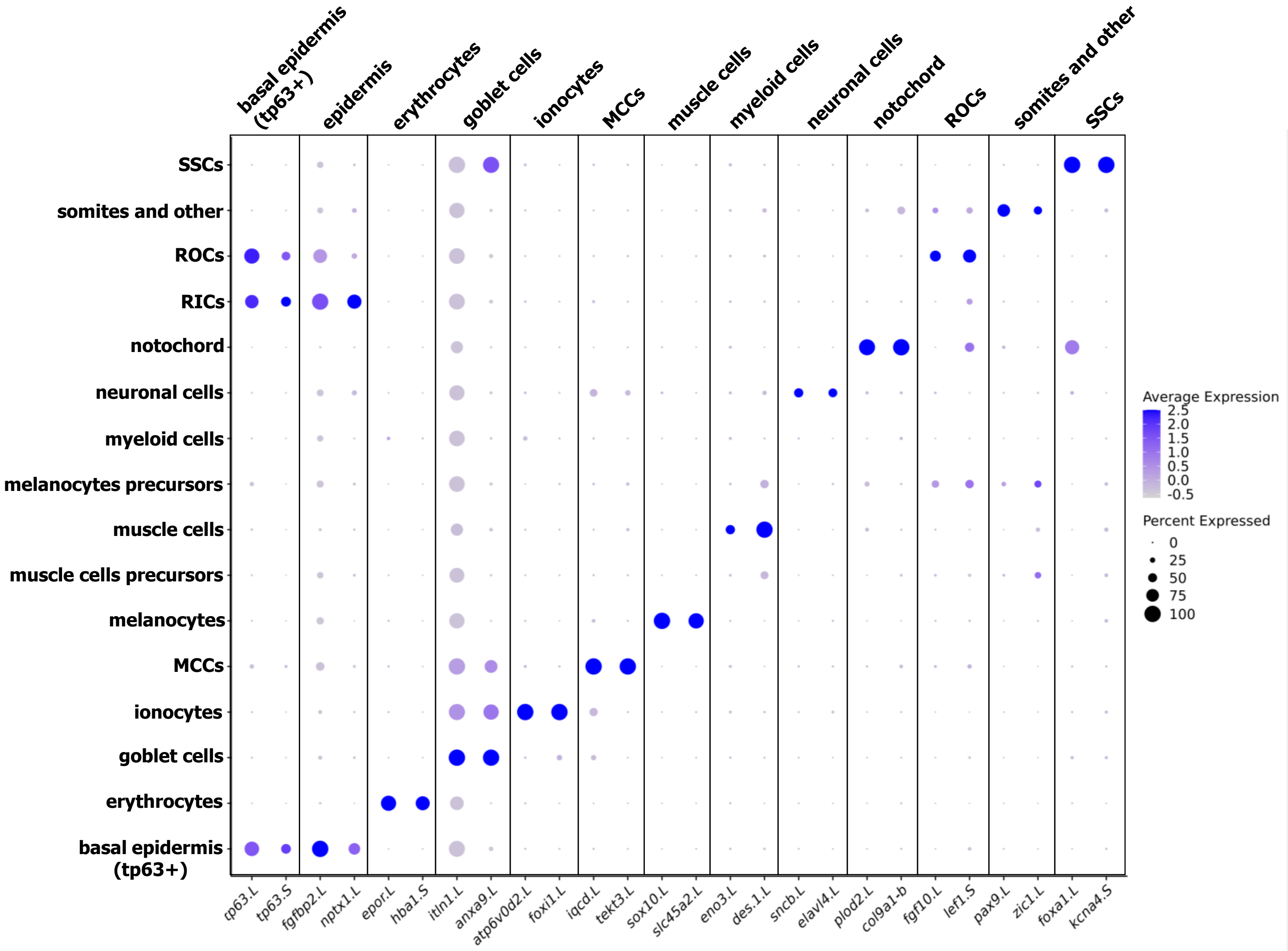


**Supplement Figure 4.** Dotplot showing the expression profile of the cell specific maker genes used to annotate the identified clusters from the scRNA-Seq experiment.

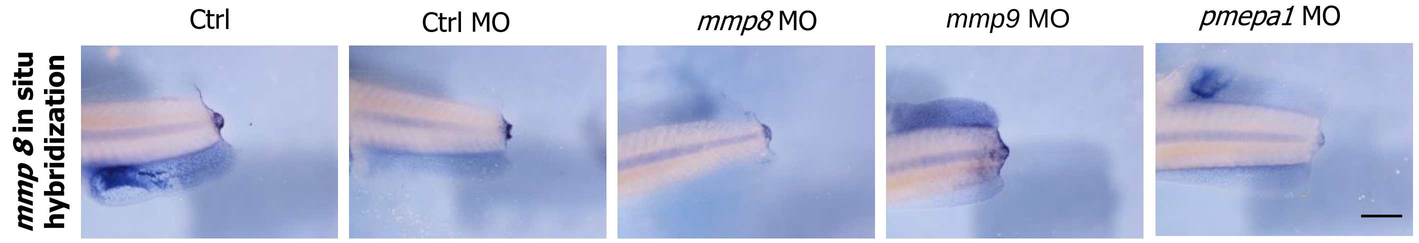


**Supplement Figure 5.** *In situ* hybridization of *mmp8* as a marker of RICs during regeneration bud formation at 1 dpa. Loss of function embryos at stage 41 showed weaker signal in the bud indicating defects in bud formation. Scale bar 500 µm.

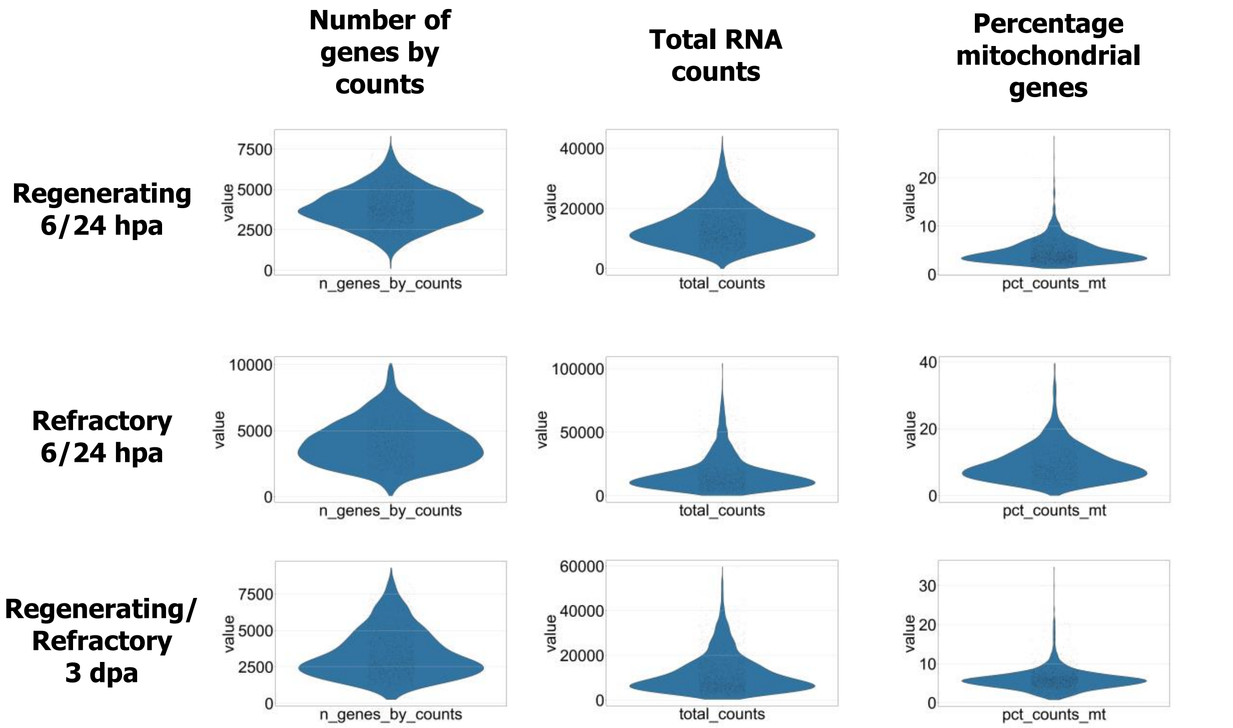


**Supplement Figure 6.** Distribution of gene counts, total RNA counts and mitochondrial genes after quality control of the spatial transcriptomic dataset.

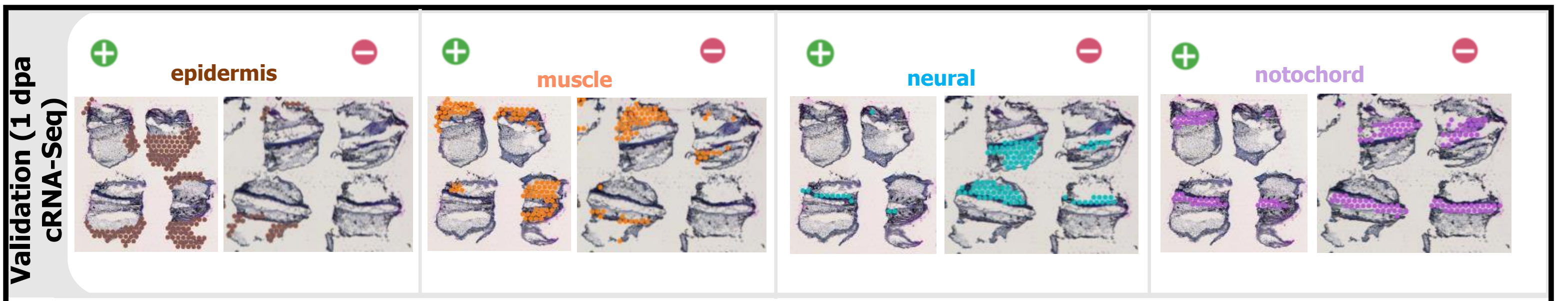


**Supplement Figure 7.** List of epidermis, muscle, neural and notochord markers from scRNA-Seq was visualized in spatial results in regenerative and refractory samples at 1 dpa.

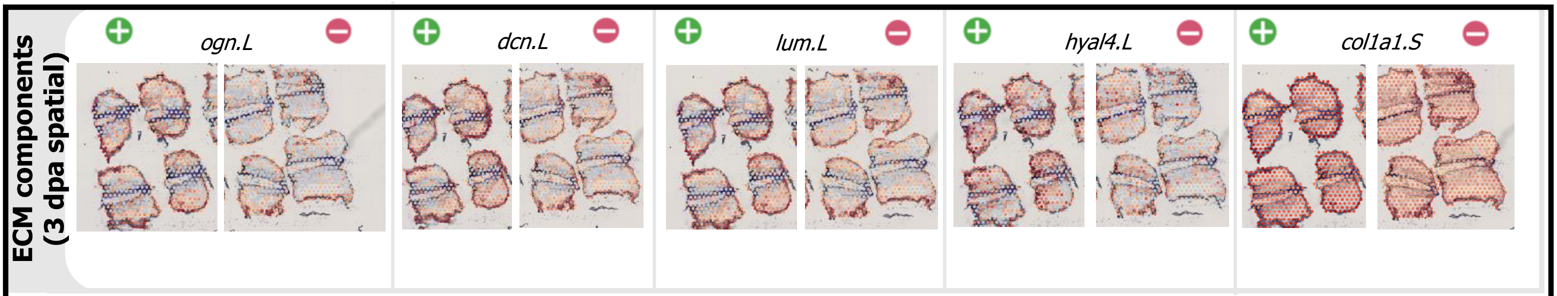


**Supplement Figure 8.** Visualization of different ECM components in regenerative and refractory samples at 3 dpa.

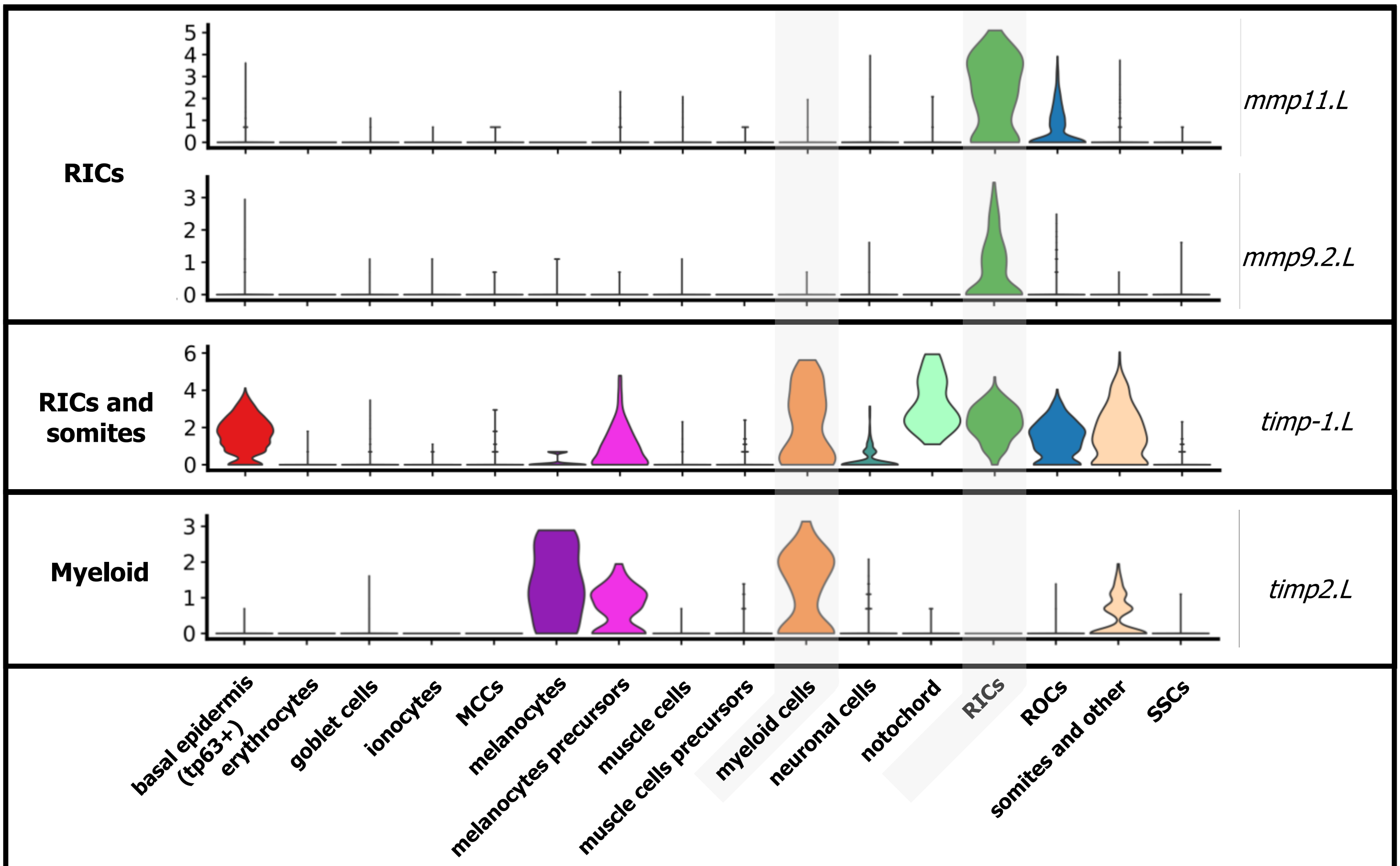


**Supplement Figure 9.** Violin plot showing selected marker genes for the cell populations identified from the scRNA-Seq experiment. Shown are other *mmp* genes that were detected as overabundant in the RICs population, *timp-1.L* in the RICs and somites, and *timp2.L* in the myeloid cells.

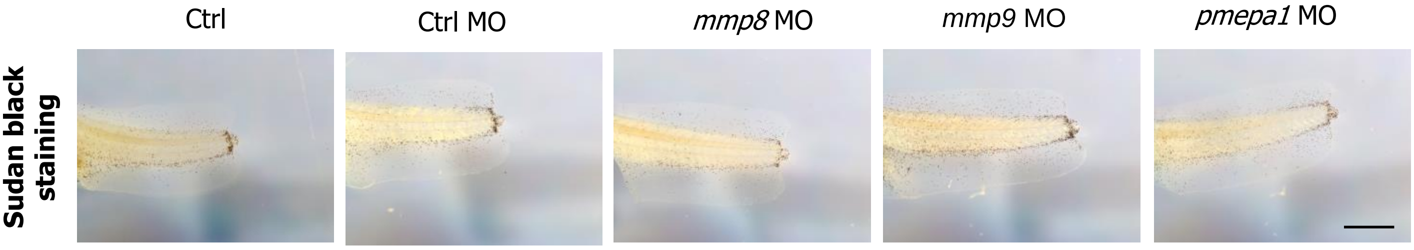


**Supplement Figure 10.** Sudan black staining at 1 dpa showing similar migration of myeloid cells in loss of function embryos at stage 41. Scale bar 500 μm.
